# Can predictive simulations of walker-assisted gait using calibrated muscle models capture subject-specific walking features in individuals with spinal cord injury?

**DOI:** 10.64898/2025.12.24.696345

**Authors:** Filippo Maceratesi, Carlos Pagès Sanchis, Josep M. Font-Llagunes, Míriam Febrer-Nafría

## Abstract

**Objective:** This study aims to evaluate whether our predictive simulation framework, coupled with different musculoskeletal model personalization methods, can reproduce the distinct subject-specific gait features of four subjects with spinal cord injury (SCI).

**Methods:** Motion capture data was collected with four SCI patients. The musculotendon parameters of the musculoskeletal models of each subject were calibrated using three different methods: one anthropometric and two functional approaches. Predictive simulations of walker-assisted gait were performed using direct collocation in an optimal control problem. The cost function included terms minimizing metabolic energy rate and muscle effort, along with additional terms reflecting the instructions of the clinicians. Post-simulation analyses were carried out to compute key gait metrics and perform inter-subject and intra-subject comparisons in both the experimental data and gait predictive simulations.

**Results:** The predictive simulations with functionally-calibrated models reproduced some distinct gait metrics for the four subjects with SCI. However, the predicted inter-subject variability of the kinematics (e.g., 7.81+/-6.04 deg for lower body joints) was generally statistically lower than the experimental one (e.g., 11.54+/-6.96 deg for lower body joints). In addition, when comparing subjects pairwise, in some cases, the predictive simulations were able to capture the similarities or discrepancies in kinematics and gait metrics between two individuals. Moreover, functionally-calibrated models yielded lower root mean square errors between the predicted and experimental lower body kinematics compared to models personalized with the anthropometric approach.

**Conclusion:** The results suggest that our predictive simulation framework can reproduce some subject-specific gait features for patients with SCI. However, further work is required to improve the realism of the musculoskeletal models (e.g., by implementing a more detailed hand-walker contact model), enhance the formulation of the predictive simulations problem (e.g., by estimating the optimal weights of the control objectives using multi-objective optimization), and include more subjects for achieving more generalizable results.

**Author summary:** Among individuals with spinal cord injury, restoring gait is a primary rehabilitation goal to improve quality of life and decrease the risk of secondary health conditions. It is fundamental to choose and tailor a specific treatment to maximize the recovery of a specific patient. Predictive simulations of gait represent a promising approach for informing these clinical decision-making processes. They would allow us to evaluate multiple “what-if” scenarios prior to a treatment and help identify the intervention with the most favorable outcome. This work serves as a building block towards a potential use of predictive simulations in clinical applications. In fact, we assess whether such simulations can reproduce and distinguish the subject-specific gait patterns of individuals with spinal cord injury. Our findings suggest that we are able to predict some key gait metrics of specific patients. However, further work is needed to improve the realism of the computational models used in the predictive simulations before such approaches can be reliably applied in clinical settings.

## Introduction

Spinal cord injury (SCI) disrupts the conduction of neural signals between the brain and the body, potentially leading to paraplegia or tetraplegia [1]. Each year, between 250,000 and 500,000 people worldwide are affected by SCI [2], often as a result of falls in the elderly or traffic accidents [3]. Among individuals with SCI, restoring gait is a primary rehabilitation goal, as impaired walking capacity reduces quality of life and increases the risk of secondary health conditions such as cardiovascular disease and hypertension. Current gait rehabilitation strategies include physical therapy (e.g., [4, 5]), overground or treadmill gait training (e.g., [6, 7]), functional electrical stimulation (e.g., [8, 9]) and robot-assisted gait training (e.g., [10, 11]). However, the response to interventions may vary significantly from patient to patient due to the level of injury, pre-existing health conditions, body composition, and overall fitness levels [12]. Therefore, it is important to choose and tailor the gait rehabilitation, e.g., by tuning the parameters of assistive devices [13, 14], to maximize the recovery of each patient.

Predictive simulations of human gait based on musculoskeletal models have great potential to support clinical decision-making processes [15]. In fact, these simulations can be used to generate a novel motion without relying on experimental movement data. Therefore, if we were able to model the effects of different treatments or assistive devices within this predictive tool, it would allow us to test various “what-if” scenarios in advance and identify the one with the greatest potential to maximize therapy outcomes for a specific patient. Most predictive tools are typically based on the assumption that the central nervous system optimizes the movement with some criteria. For this reason, algorithms using a black box neural controller with optimization or reward policies are often employed to predict human motion [16]. Recent studies have used reinforcement learning (e.g., [17, 18]), multiple shooting (e.g., [19, 20]) and direct collocation (e.g., [21, 22]) techniques to perform predictive simulations.

In several studies, the impaired gait of subjects across different clinical populations was predicted to either elucidate the causes-effect relationships of gait impairments, simulate rehabilitation procedures, model assistive devices, or predict surgical outcomes. For example, some researchers [23–25] predicted gait adaptations in osteoarthritis patients to modify external knee adduction torques, while Reinbolt *et al.* [26] also estimated their functional recovery after high tibial osteotomy surgery. Other researchers used predictive simulations to investigate post-stroke gait, by developing a synergy-controlled neuromusculoskeletal model [27], assessing how gait impairments and asymmetries affect metabolic cost [28], and evaluating personalized functional electrical stimulation training protocols [29]. Falisse *et al.* [30] analyzed the effects of musculoskeletal and motor control impairments on the walking pattern of a specific child with cerebral palsy. This work was recently expanded to include more children and generalize the results [31]. Predictive simulations were also applied in amputees, by testing how transtibial gait is influenced by prosthetic foot characteristics [32] and assessing the ability of direct collocation techniques to reproduce pathological gait with a lower-leg prosthesis [22]. Eskinazi and Fregly [33] applied their novel framework for performing concurrent muscle, joint contact, and joint kinematic simulations to replicate the gait of a patient with total knee replacement. For individuals with SCI, simulation frameworks were used to personalize knee trajectory parameters and optimize ankle stiffness in knee–ankle–foot orthoses for assisted gait [34, 35]. A major challenge faced in this line of work is that the predictive simulations are generally performed for a single patient only. As a result, even if a framework is validated for that individual, it may not replicate the walking motion of other patients. This challenge becomes even more pronounced when predicting gait with an assistive device, as the interaction between the device and the user can vary substantially across individuals. Consequently, it remains unclear whether such models are capturing a generic impaired gait that broadly reflects the walking patterns of a given population, or truly reproducing subject-specific gait features, such as individual compensatory strategies.

To employ predictive simulation tools in clinical applications, it is fundamental that the musculoskeletal models used in such simulations represent as closely as possible the studied patients. For this reason, high levels of complexity and personalization are needed when developing such musculoskeletal models. The most widely used approach to actuate these models in gait predictive simulations is using Hill-type muscle models in the lower body and ideal torques in the upper body. Ideal torques simply allow the model to generate the amounts of resultant joint moments necessary to perform a certain motion. Hill-type models represent single musculotendon units, and they generate forces through a contractile, passive elastic, and tendon elastic element [36]. The parameters of the Hill-type muscle actuators used in these musculoskeletal models can be calibrated to match the muscular properties of the studied individuals. Two overarching methods are employed in literature to calibrate these parameters: anthropometric and functional approaches. In the former method, the musculotendon parameters are scaled based on the skeletal dimensions of the specific subject, while preserving the operating conditions of the muscles in a reference model [37, 38]. This approach is used to scale the optimal fiber lengths (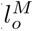) and the tendon slack length (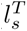) of the muscle models. On the other hand, functional approaches consist in optimizing the musculotendon parameters, while minimizing the difference between experimental and model-based joint torques [39–43]. Nearly all studies using a functional approach calibrate 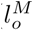 and 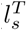, with some also adjusting other parameters related to contraction dynamics (e.g., maximum isometric force, optimal pennation angle) and activation dynamics (e.g., activation and deactivation time constants) [44–46]. It is not clear whether using simple anthropometric approaches is sufficient or functional methods that rely on experimental data are necessary to improve the results of the gait predictions [47]. In addition, more work is required to identify which musculotendon parameters are important to calibrate for predictive simulations in patients with SCI. In fact, even though muscle force estimation is primarily sensitive to the 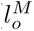 and 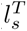 values [48–52], calibrating the parameters that govern muscle activation dynamics might be fundamental to reproduce the delayed and reduced neuromuscular responses characteristic of SCI patients.

Therefore, the overall aim of this study is to assess whether our predictive simulation framework [47], combined with different musculoskeletal model personalization approaches, can capture and distinguish the subject-specific gait features of four SCI patients, rather than predicting a generic assisted gait. To achieve this goal, we structured the study in three parts. First, the experimental data were analyzed to characterize the gait of each subject and quantify inter-subject differences in walking patterns. Second, we compared the predictive simulations of models with functionally-calibrated 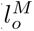 and 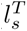 values against the experimental measurements. Third, we examined how musculoskeletal models with three different levels of personalization influenced the gait prediction results: an anthropometrically-scaled model, the aforementioned functionally-calibrated model, and an extended functionally-calibrated model with personalized activation time constants. In all three parts of the study, we analyzed both lower and upper body kinematics, as well as gait metrics that are commonly reported in clinical studies involving patients with SCI.

## Materials and methods

### Ethics statement

The experimental protocol was approved by the Ethical Committee of the Complejo Hospitalario Universitario de Toledo (approval date: November 30, 2022; ID code: 929). The recruitment of the subjects began on January 9, 2023 and ended on July 26, 2024. All participants gave written consent to participate in the experiments.

### Experimental measurements with SCI subjects

The experimental activities were carried out in the biomechanics laboratory of the Hospital Nacional de Parapléjicos (Toledo, Spain). Four individuals with SCI participated in the experiments (see Table 1 for more information about each subject). They were instrumented with 43 reflective markers, corresponding to an extended version of the PlugInGait Fullbody protocol developed by Vicon, and 8 EMG electrodes (Cometa WavePlus, Cometa srl, Milano, Italy) per leg to record the activity of the following muscles: tibialis anterior, soleus, gastrocnemius medialis, vastus lateralis, rectus femoris, biceps femoris, semitendinosus and gluteus medius. The electrodes were placed following the guidelines of Surface ElectroMyoGraphy for the Non-Invasive Assessment of Muscles (SENIAM) [53]. Following the subjects’ preparation, maximum isometric tests were performed for the hip abductors, knee flexors, knee extensors, ankle dorsiflexors and ankle plantar flexors. Each muscle group was tested three times, with a one-minute rest between measurements. Subsequently, a motion capture system (Vero 2.2, Vicon Motion Systems Ltd., Oxford, Oxfordshire, UK) and ground-embedded force plates (Kistler 9286AA, Kistler Group, Winterthur, Switzerland) were used to record the markers’ trajectories and the ground reaction loads during two activities: gait and sit-to-stand-to-sit. These two functional tasks were selected as they were the only ones the subjects were able to perform from the set of motions used in similar modeling studies [42]. In addition, a static trial was recorded to measure the weight of the individuals and for scaling the musculoskeletal models. All participants were assisted by a walker during the motion capture experiments.

**Table 1.** Information about the subjects with SCI (S1-S4) that participated in the experiments.

|  | **S1** | **S2** | **S3** | **S4** |
| --- | --- | --- | --- | --- |
| **Age** | 57 | 67 | 40 | 61 |
| **Height/weight** | 1.76 m/80 kg | 1.58 m/52 kg | 1.85 m/94 kg | 1.85 m/107 kg |
| **WISCI-II** | 16 | 9 | 16 | 13 |
| **ASIA** | C | D | C | D |
| **10MWT** | 6.39 s | 47.23 s | 18.78 s | 28.58 s |
| **TUG** | 14.61 s | 57.19 s | 27.99 s | 42.49 s |

The Walking Index for Spinal Cord Injury II (WISCI-II) measures the type of assistance required by the individual to walk, ranging from 0 (unable to stand) to 20 (walking with no device and no assistance).
The American Spinal Injury Association (ASIA) impairment score classifies the severity of the injury, ranging from A (complete injury with no preserved function) to E (normal function).
The 10 Meter Walk Test (10MWT) is a gait performance assessment used to measure the time required to walk 10 meters at a self-selected speed.
The Timed Up and Go (TUG) test measures the time required by the subject to stand from a chair, walk 3 meters, turn, walk back and sit down on the chair.

Following the experimental procedures, the raw measurements were processed to prepare them for use in MATLAB and OpenSim. Marker trajectories were interpolated to fill any gaps in the recordings. The ground reaction forces were filtered using a 4th-order zero-phase-lag Butterworth filter with a cut-off frequency of 6 Hz, in accordance with several previous studies (e.g., [41, 44]). EMG signals were also processed following the approach described in [42]. First, they were band-pass filtered between 20–400 Hz, followed by full-wave rectification to convert negative amplitudes to positive values. Second, a 2nd-order Butterworth low-pass filter with a 10 Hz cut-off was applied. Finally, EMG signals for each muscle were normalized to the peak value observed during the entire set of experiments. An additional normalization step was performed during the calibration of the musculotendon parameters (see subsection on muscle calibration problem formulation), since achieving true maximal contractions is challenging in practice [54]. Because some muscles shared innervation and contributed to the same functional actuation, they were assigned identical EMG patterns. As a result, the 16 recorded EMG profiles were attributed to 36 musculotendon units in the model (see S1 Fig in S1 File).

### Musculoskeletal modeling

#### OpenSim analyses

A generic, full-body OpenSim model was employed for this study [55]. The model had 37 degrees of freedom (DOFs; 14 in the lower limbs, 6 in the pelvis, 3 in the lumbar joint, and 14 in the upper limbs) and 40 Hill-type muscle actuators per leg. The musculoskeletal system was scaled to match the anthropometric dimensions of the four subjects using the “Scale Tool” in OpenSim. This process linearly scaled the 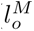 and 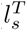 values of each muscle based on the change in musculotendon lengths after scaling. Moreover, inverse kinematics (IK) and inverse dynamics (ID) analyses were carried out on OpenSim to compute the reference data for the calibration problems and the post-simulation analyses. The marker trajectories were input into the IK tool to obtain the musculotendon lengths, moment arms, generalized coordinates, velocities and accelerations. The joint torques and residual loads were calculated through ID.

#### Muscle modeling and calibration

The muscle activation dynamics reflected the relationship between muscle excitation *e^M^* and activation *a^M^*with a first-order differential equation [36]:

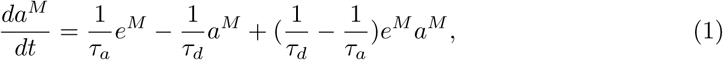

where *τ_a_* and *τ_d_* are the activation and deactivation time constants, respectively. These two parameters tune how quickly a muscle can produce and release force in response to neural excitations. Furthermore, as mentioned above, the contraction dynamics was defined by Hill-type models, characterized by active force-length (**f***^l^*), passive force-length (**f***^P^ ^E^*), active force-velocity (**f***^V^*) and tendon force-length (**f***^T^*) dimensionless curves:

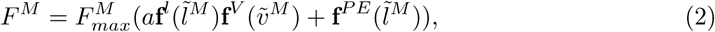

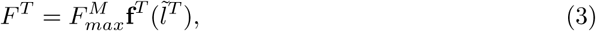

where *F^M^* and *F^T^* are the muscle and tendon forces, respectively. The 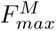 parameters for each muscle were computed using regression equations that relate muscle volume to each subject’s mass and height [56], and subsequently linearly scaled according to the lower extremity motor scores (LEMS) of the patients. LEMS are clinical scales performed by clinicians to score the strength of lower extremity muscle groups of SCI subjects [57].

The Hill-type musculotendon parameters were calibrated using three different approaches to generate three models with varying levels of personalization per subject. The first method involved non-linearly scaling the 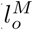 and 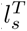 values using the anthropometric method developed by Modenese *et al.* [38]. This approach consists in mapping the normalized muscle operating conditions of an existing reference model (i.e., the generic model described in the previous section), onto a scaled model of different anthropometric dimensions for equivalent joint configurations. For the second approach, we used the baseline formulation of our functional calibration framework to estimate musculotendon parameters described in [47]. To summarize this method, a multi-phase optimal control problem (OCP) was solved on GPOPS-II [58] with direct collocation. ADiGator [59], a MATLAB automatic differentiation package, was used to improve the computational efficiency of the problem. Three parameters of 18 muscles per leg (specified in S1 Fig in S1 File) were estimated through this OCP: 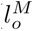, 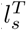 and a factor*k_EMG_* that was used to scale the EMG data as a further normalization of the signals. The calibration in this study differs from the one described in our previous work in two main ways: (i) the solution space of 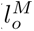 and 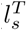 was reduced to *±*10% of the non-linearly scaled values to avoid excessively high passive forces during gait, and (ii) only gait and sit-to-stand-to-sit data were used in the OCP, as the patients were unable to perform the other functional motions included in our previous study. The third approach consisted in calibrating the *τ_a_* parameter of the same 18 muscles per leg through a similar functional method, using optimal control. *τ_a_* was constrained to vary between 5 ms and 35 ms, while the value of 15 ms was used as both the initial guess of the calibrated muscles and the generic value for uncalibrated muscles [44]. *τ_d_* was imposed to be proportional to *τ_a_*such that *τ_d_*= 4*τ_a_*, based on Zajac’s model [36]. It is important to note that the only parameter that was allowed to vary in the OCP was *τ_a_*, while 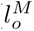, 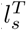 and *k_EMG_* were kept identical to the values obtained with the second approach. This allowed for a more direct comparison between the predictive simulations of the models calibrated using the second and third calibration methods. In fact, through this comparison, we could clearly examine the effects of the calibration of *τ_a_*. Henceforth, musculoskeletal models with musculotendon parameters calibrated using the first approach are referred to as non-linearly scaled models (NSMs). Musculoskeletal models calibrated using the second and third approaches are referred to as functionally calibrated models (FCMs and FCMs-*τ_a_*, respectively).

#### Contact modeling

The foot-ground contact model comprised 16 spring-damper units on each foot. The normal force in each element was generated using a linear spring with non-linear damping [60], and the tangential force in each element was calculated using a simple continuous friction model [60]. A sensitivity analysis in a previous study demonstrated that changing the parameters of the foot-ground contact model did not significantly influence predictive simulations of walking [22]. For this reason, we used parameters that were previously calibrated for a young healthy individual [61] and linearly scaled them according to the body weight of the subjects with SCI.

The hand-walker interaction was modeled through a set of constraints, defined in the OCP of the predictive simulation framework (described in the next section). Sliding kinematic constraints were defined in the anteroposterior direction to keep the centers of the hands within a certain distance from each other and the ground, corresponding to the width and height of the walker. The interaction with the walker was modeled by applying three external forces at each hand, which were included as controls in the OCP. In addition, the total vertical hand reaction force (HRF) was constrained, within a tolerance, to match the percentage of weight supported by the walker (PWS). This percentage was defined assuming that during unassisted slow gait the maximum vertical ground reaction forces equal body weight [62]. Accordingly, for SCI individuals walking with a walker, PWS was computed from the mean peak vertical ground reaction forces of the feet across trials as the remaining force required to support body weight. The estimation of the hand reaction forces through optimal control relies on the fulfillment of dynamic consistency, by minimizing the residuals of the pelvis.

### Predictive simulations of assisted gait

An OCP was developed on GPOPS-II and solved with direct collocation to perform predictive simulations of one assisted-gait cycle. The states of the problem included: the joint coordinates, joint velocities, muscle activations in the lower body, normalized tendon forces and activations in the upper body for the ideal torques. The controls of the problem included: joint accelerations, time derivative of muscle activations in the lower body, ground reaction loads, time derivative of normalized tendon forces, hand reaction forces and upper body excitations. Three different initial guesses were tested for the predictions with FCMs: (i) the experimental gait of the SCI participants (used for the corresponding subject with SCI), (ii) the experimental gait data of a healthy participant, and (iii) the mean solution obtained from the predictive simulations using the first initial guess across all SCI subjects. The differences between the solutions with these initial guesses were either comparable or significantly smaller than the experimental intra-subject variability (see S2 Fig in S1 File). Therefore, for the remaining models, only the first initial guess was used. The duration of the simulation was bounded to be between 80% and 120% of the mean duration of the gait cycles in the experiments. The static parameters of the problem comprised the beginning and end of the swing phase of each leg, expressed as percentages of the gait cycle. These parameters were bounded to remain within *±*10% of the experimental values. The initial guess for these parameters corresponded to the experimental data.

The cost function *J* was adapted from a similar predictive simulation framework [22], with some additional terms accounting for the clinicians’ instructions provided to the SCI patients during gait training:

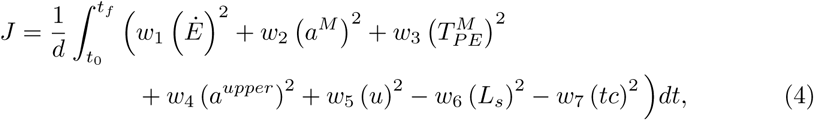

where, *d* is the anteroposterior distance traveled by the pelvis in the predicted gait, *t*_0_ and *t_f_* are the initial and final time of the simulation, respectively, Ė is the metabolic energy rate calculated using the phenomenological model described in [63], *a^M^* is, as previously defined, the muscle activations, 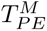 is the passive joint torques computed through the passive muscle forces, *a^upper^* is the activations in the upper body for the ideal torques, *u* is the controls of the OCP which are minimized for regularization purposes, *L_s_* is the stride length of the feet, *tc* is the toe clearance of both feet from the ground at the beginning of the swing phase, and *w*_1−7_ are the weight factors attributed to each cost function term. Among the cost terms, *L_s_* and *tc* were maximized to reflect the instructions of the clinicians directing patients to take the largest possible steps and increase toe clearance to improve gait and enhance safety.

The skeletal dynamics was implemented implicitly by ensuring that the residual loads remained low through the path constraints of the OCP [32]. Additional path constraints enforced the equilibrium between the muscle and tendon forces to model the Hill-type contraction dynamics implicitly, following the formulation described in [64]. The activation dynamics of the muscles was expressed as a set of linear equality and inequality constraints, as explained in [65]. This allowed us to remove muscle excitations from the design variables of the OCP and improve its computational efficiency. Moreover, as previously mentioned, path constraints were used to limit the positions of the hands and the HRFs (see the subsection regarding the walker modeling for more details). The entire set of path constraints is shown and explained in the supplementary material (see S1 File). Event constraints were included to bound the stride length to the experimentally measured value (within a tolerance) and to enforce kinematic periodicity over the gait cycle (also within a tolerance). The dynamic constraints were simply defined as the derivative relationships of the states.

### Post-simulation analyses

Key gait metrics were computed in all gait cycles from both the experimental measurements and the results of the predictive simulations: the circumduction, circuity index, mean trunk inclination, muscles co-contractions and the range of motions (ROMs) in the lower and upper body. The circumduction was estimated as the maximum lateral difference of the ankle marker between stance and swing phase. The circumduction of both feet were computed and averaged together. The circuity index was calculated as the ratio between the distance covered by the centre of mass in the transverse plane and its euclidean distance between the start and end point of each gait cycle. The mean trunk inclination was defined as the average of the addition of the lumbar flexion and the forward pelvic tilt. The co-contractions were computed for each antagonistic muscle pair measured with EMGs, using the method described in [66]. The ROMs were calculated separately for the lower and upper body, excluding the DOFs at the pelvis. Other important gait metrics, such as the stride length, gait speed, stance/swing durations and maximum vertical ground reaction forces were omitted from the analysis since they were bounded in the OCP of the predictive simulations to match the experimental values (within a tolerance).

We performed an inter-subject and intra-subject analysis of the experimental kinematics and gait metrics. For the kinematics, we computed the inter- and intra-subject variability and correlation; whereas, for the gait metrics, we only calculated the inter- and intra-subject variability. The inter-subject variability of the kinematics was quantified by computing the root mean square error (RMSE) between the joint angles of every pair of subjects. On the other hand, the inter-subject correlation of the kinematics was quantified by computing the coefficient of determination (R^2^) between the joint angles of every pair of subjects. The intra-subject variability and correlation of the kinematics was estimated using the same metrics, but applied to all pairs of gait cycles measured for each subject. For the gait metrics, the inter- and intra-subject variability was calculated using RMSEs in the case of the muscles co-contractions (since it is a continuous variable) and simple differences in case of the other metrics. Then, the inter-subject and intra-subject variability/correlation were compared against each other to ensure that the subjects showed sufficiently distinct walking patterns, allowing us to later assess whether the predictive simulations could capture and differentiate these subject-specific gaits.

The inter-subject variability and correlation were also computed for the predictive simulations with FCMs, using the same method described above. These quantities were then compared to the experimental intra-subject variability and correlation to verify whether the predicted gait patterns of different subjects were distinct. We also compared the predicted and experimental inter-subject variability/correlation to assess if they were similar or not significantly different. Moreover, the variability and correlation between the predictive simulation results and the experimental measurements of the corresponding subject were estimated. These quantities were compared to the experimental and predicted inter-subject variability/correlation to verify whether the predictions were capturing subject-specific gaits. Furthermore, we assessed whether each simulation was predicting the kinematics and gait metrics of the experimental measurements of the corresponding subject more accurately than the experimental data other subjects. Lastly, to determine which type of model predicted more accurately the gait pattern of the corresponding subject, we compared the RMSE and R^2^ values (for the kinematics) and the differences and RMSE (for the gait metrics) between the experimental data and the simulation results of NSMs, FCMs and FCMs-*τ_a_*.

All quantitative comparisons are summarized in Table 2. For each of these comparisons, we performed a statistical analysis to examine whether there were significant differences. To do so, we first checked if the data samples being compared were normal using the Lilliefors test. If both datasets being compared were normal, the t-test was employed to perform the statistical analysis; whereas if either dataset was not normal, the Mann-Whitney U-test was used. Statistical significance was achieved when a p-value was less than 0.05. Moreover, qualitative analyses of the following key joint angles were carried out for both the experimental data and the predictive simulation results: hip flexion/extension (hipFE), hip adduction/abduction (hipAA), hip internal/external rotation (hipIE), knee flexion/extension (kneeFE) and ankle plantar flexion/dorsiflexion (anklePD).

**Table 2.**
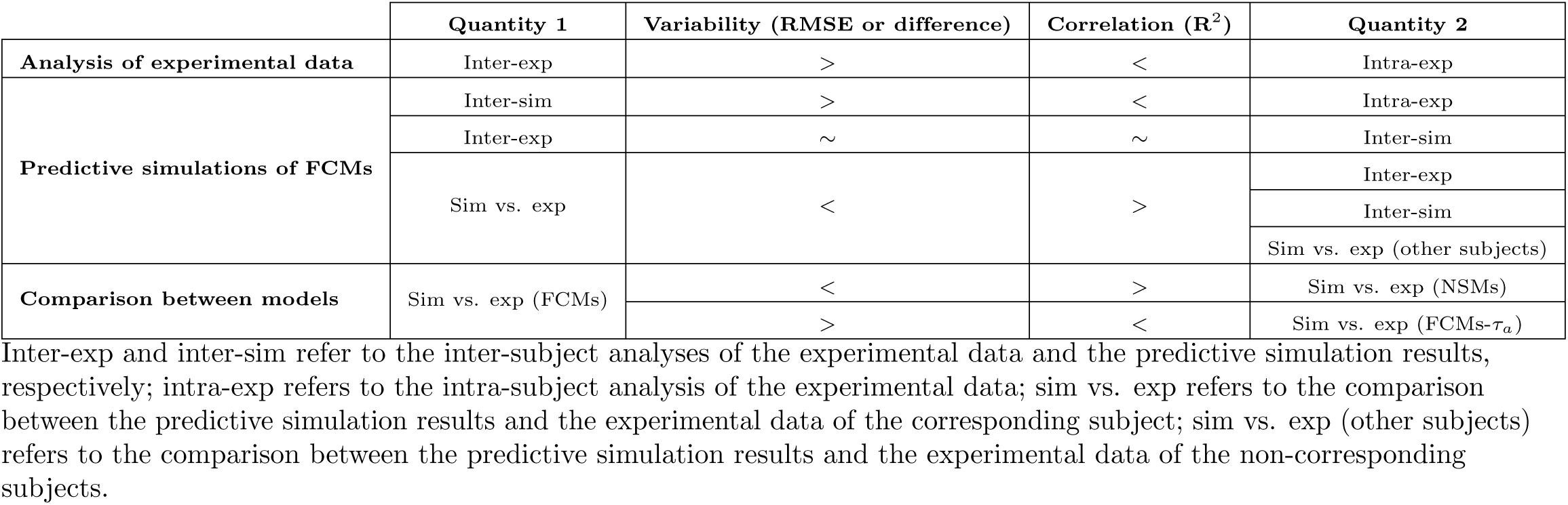
Summary of the post-simulation quantitative analyses. The third and fourth columns report the desired outcomes of the quantity 1-quantity 2 comparisons in the analysis of the experimental data and the predictive simulations of FCMs, as well as the expected outcomes in the comparison between the models.

| | Quantity 1 | Variability (RMSE or difference) | Correlation ($R^2$) | Quantity 2 |
| --- | --- | --- | --- | --- |
| Analysis of experimental data | Inter-exp | > | < | Intra-exp |
| Predictive simulations of FCMs | Inter-sim | > | < | Intra-exp |
|  | Inter-exp | ~ | ~ | Inter-sim |
|  | Sim vs. exp | < | > | Inter-exp |
|  |  |  |  | Inter-sim |
|  |  |  |  | Sim vs. exp (other subjects) |
| Comparison between models | Sim vs. exp (FCMs) | < | > | Sim vs. exp (NSMs) |
| | | > | < | Sim vs. exp (FCMs-$\tau_a$) |

Inter-exp and inter-sim refer to the inter-subject analyses of the experimental data and the predictive simulation results, respectively; intra-exp refers to the intra-subject analysis of the experimental data; sim vs. exp refers to the comparison between the predictive simulation results and the experimental data of the corresponding subject; sim vs. exp (other subjects) refers to the comparison between the predictive simulation results and the experimental data of the non-corresponding subjects.

## Results

### Analysis of experimental data

The subjects with the highest WISCI-II score (S1 and S3) showed good bilateral symmetry and high consistency across gait cycles (Fig 1). S2 exhibited limited hip extension during the stance phase, indicating muscle weakness that prevented longer step lengths. S2 also showed a reduced knee flexion during the swing phase of both legs, which was compensated for by a shorter swing phase and by some circumduction movement of the leg. S4 walked with a moderately high asymmetry between right and left legs, which was especially noticeable in the hipFE and kneeFE angles. The left knee of S4 remained slightly flexed throughout the entire gait cycle and showed limited flexion during swing. When comparing subject pairwise (S1 Table in S1 File), the RMSE between the experimental lower body kinematics of S2 and S4 was low compared to other subject pairs. In contrast, the largest RMSE in experimental lower body kinematics was observed between S2 and S3. For the upper body, the mean R^2^ between the experimental kinematics of S1 and S3 was higher than in other subject pairs. Furthermore, the joints ROMs in the upper body of S2 were substantially higher than the other subjects (Fig 2). This increased upper body mobility of S2 caused large COM displacements along the mediolateral direction, thereby increasing the circuity index. The mean muscles co-contraction of S3 was lower than other subjects, primarily due to the minimal co-contractions between the hamstrings and quadriceps in both legs. When comparing the gait metrics of each subject (S2 Table in S1 File), S1 and S4 showed similar circuity indices, whereas S1 and S2 had similar mean trunk inclinations.

**Fig 1.**
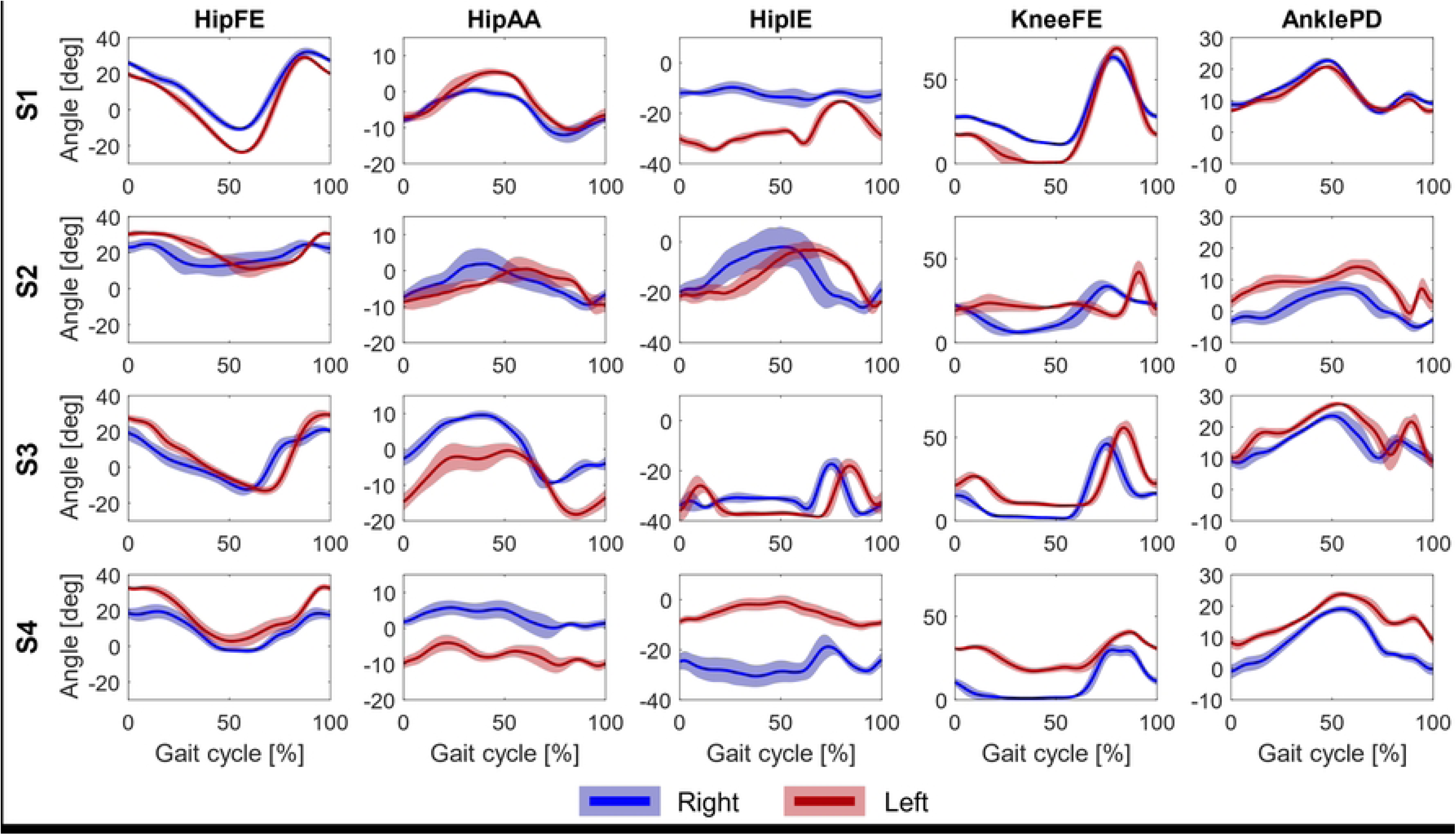
Experimental joint angles profiles. The experimental mean angle profiles (lines) and the standard deviations (shaded areas) of hipFE, hipAA, hipIE, kneeFE and anklePD of the four subjects.

**Fig 2.**
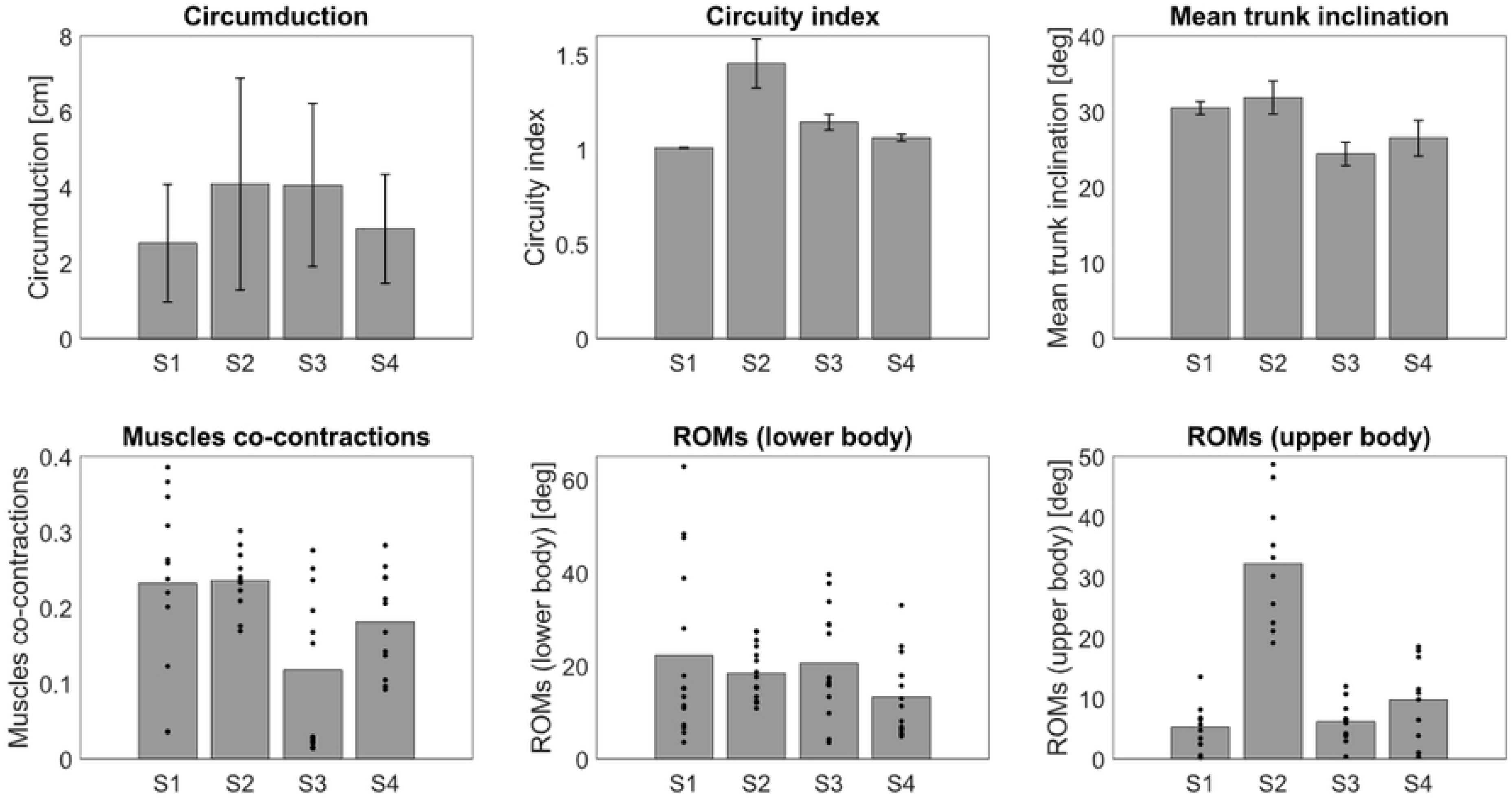
Experimental gait metrics. The top row includes the mean experimental circumduction, cicuity index and mean trunk inclination values (bars), with their standard deviations (error bars) of the four subjects. The bottom row includes the mean experimental values of muscles co-contractions, ROMs in the lower and upper body (bars), along with the mean value of each muscle pair and DOF (dots) for co-contractions and ROMs, respectively.

The inter-subject analysis showed statistically significant higher RMSE and lower R^2^ values than the intra-subject analysis in both the lower and upper body kinematics, indicating greater variability and weaker correlations across different subjects (Fig 3). These findings highlight that each subject walked in their own unique way, allowing us to assess whether the predictive simulations can capture these individual variations. In addition, among all subjects, S2 (the individual with the lowest WISCI-II score) exhibited the highest intra-subject variability, indicating inconsistencies between gait cycles. Furthermore, for most of the gait metrics, the experimental inter-subject variability was significantly higher than the experimental intra-subject variability (Fig 4). The only exceptions occurred in circumduction, where for S2 and S3 no statistically significant differences was observed between intra- and inter-subject variability.

**Fig 3.**
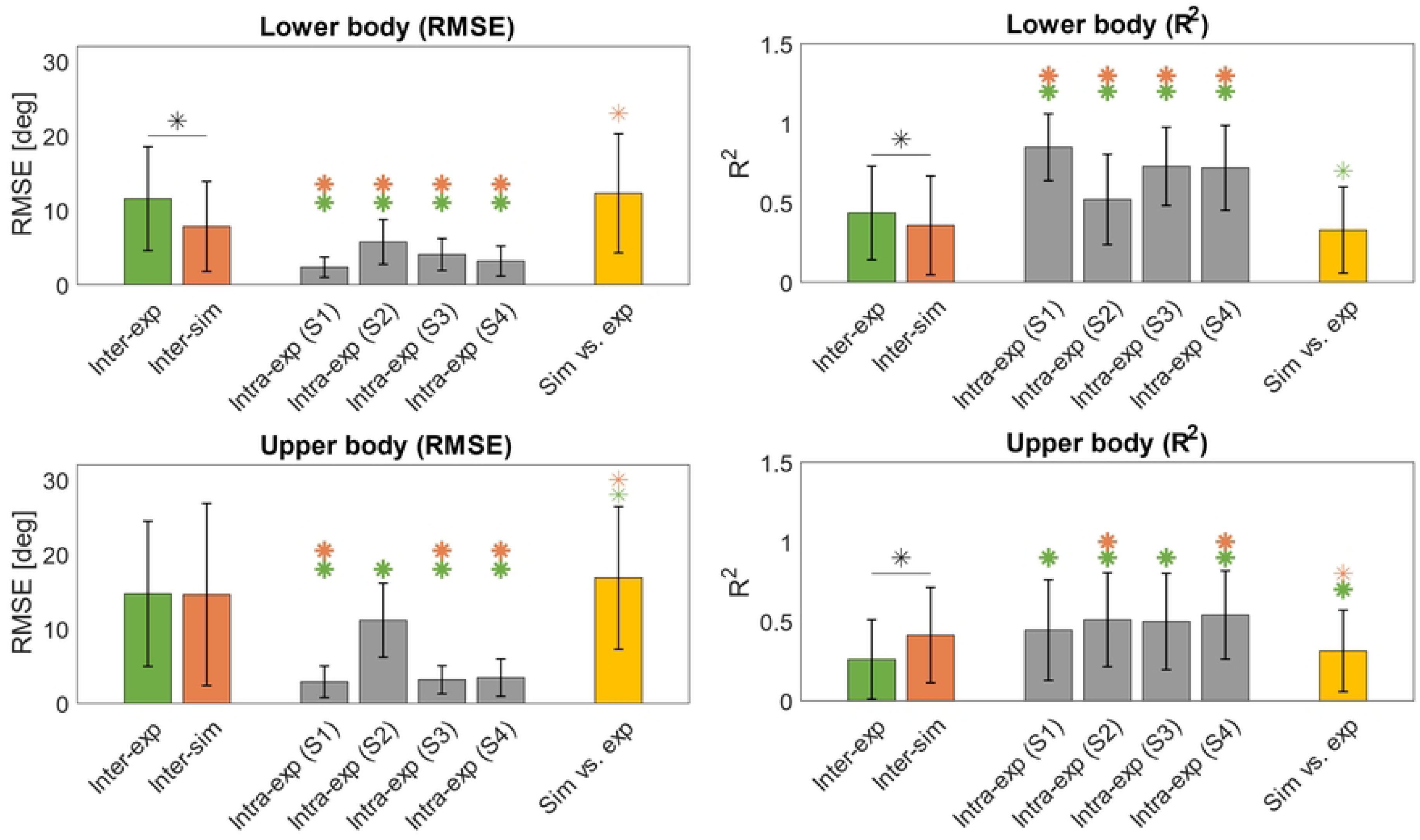
Inter-subject and intra-subject analyses, along with the comparison between experimental data and simulation results of the FCMs, for the lower and upper body kinematics. The bar plots indicate the mean values, whereas the error bars indicate the respective standard deviations. Asterisks indicate statistically significant difference. If the asterisk is bold, the statistically significant difference represents a desired outcome (as specified in Table 2).

**Fig 4.**
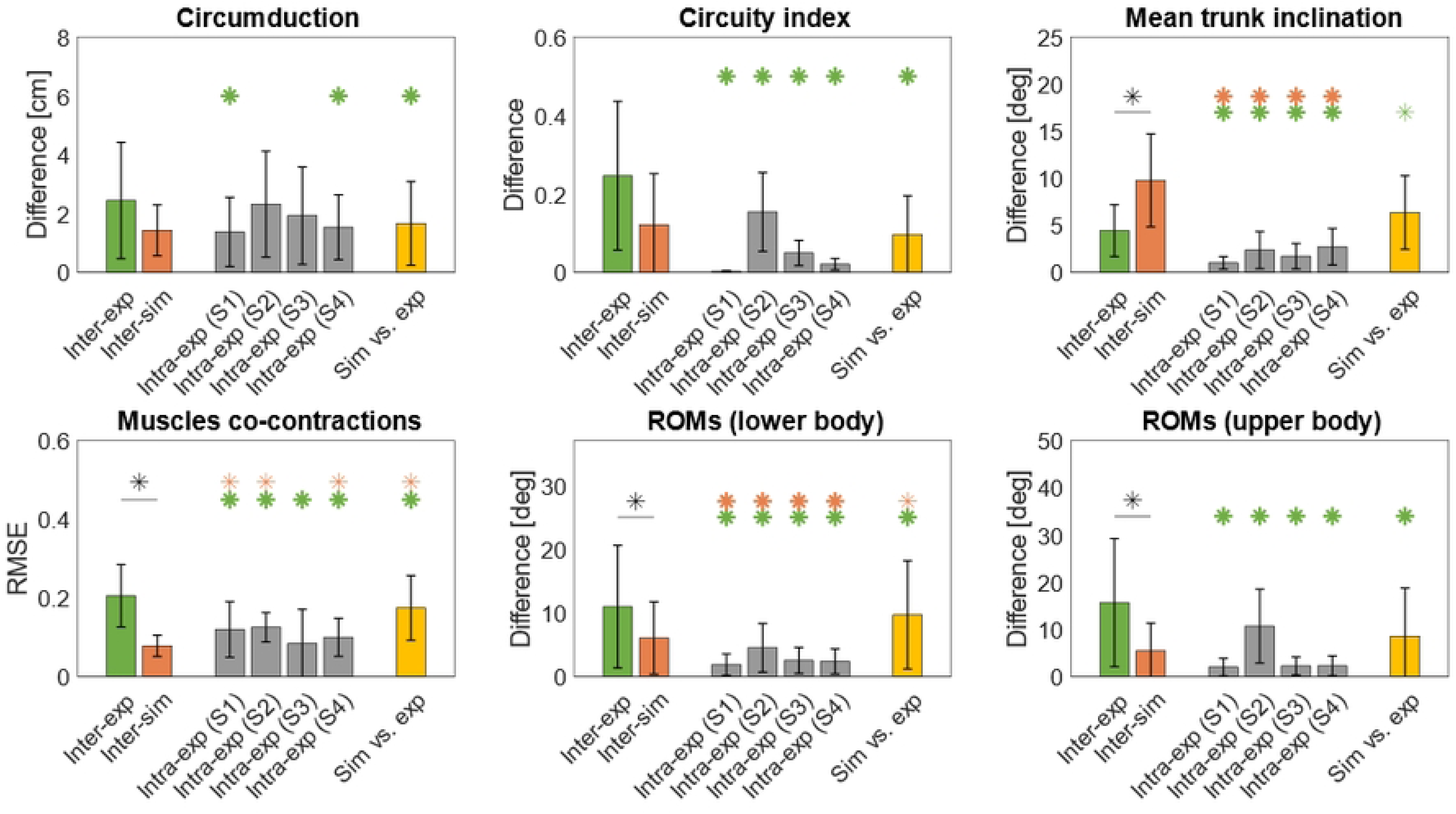
Inter-subject and intra-subject analyses, along with the comparison between experimental data and simulation results of the FCMs, for six gait metrics. The bar plots indicate the mean values, whereas the error bars indicate the respective standard deviations. Asterisks indicate statistically significant difference. If the asterisk is bold, the statistically significant difference represents a desired outcome (as specified in Table 2).

### Predictive simulations of FCMs

The predicted anklePD angle profiles (Fig 5) have some similarities with the corresponding experimental profiles (Fig 1). In fact, the left feet of S1 and S2 struck the ground in the same way in both the simulations and the experiments. S1 touched the ground with the heel (i.e., dorsiflexing the ankle) and S2 landed on the ground with a flat-foot. However, we erroneously predicted that S1 would touch the ground with a right flat-foot. The predicted ROMs of the right and left anklePD angles of S3 matched fairly well the experimental ones. In addition, the predictions of S3 correctly captured the bi-lateral symmetry in the ankle joints. Regarding the kneeFE angles, the simulations consistently predicted stiff knees in S1, S3 and S4. Even though the left knee of S4 was also stiff throughout the experimental gait cycles, this did not occur for S1 and S3. The peak knee flexion values were better predicted in the left legs of S3 and S4, compared to the other subjects. Moreover, for all subjects, the predicted hipAA and hipIE angles profiles remained much more constant compared to the experimental data. The ROM of the hipFE angle was accurately predicted in both legs of S2 and in the right limb of S3.

**Fig 5.**
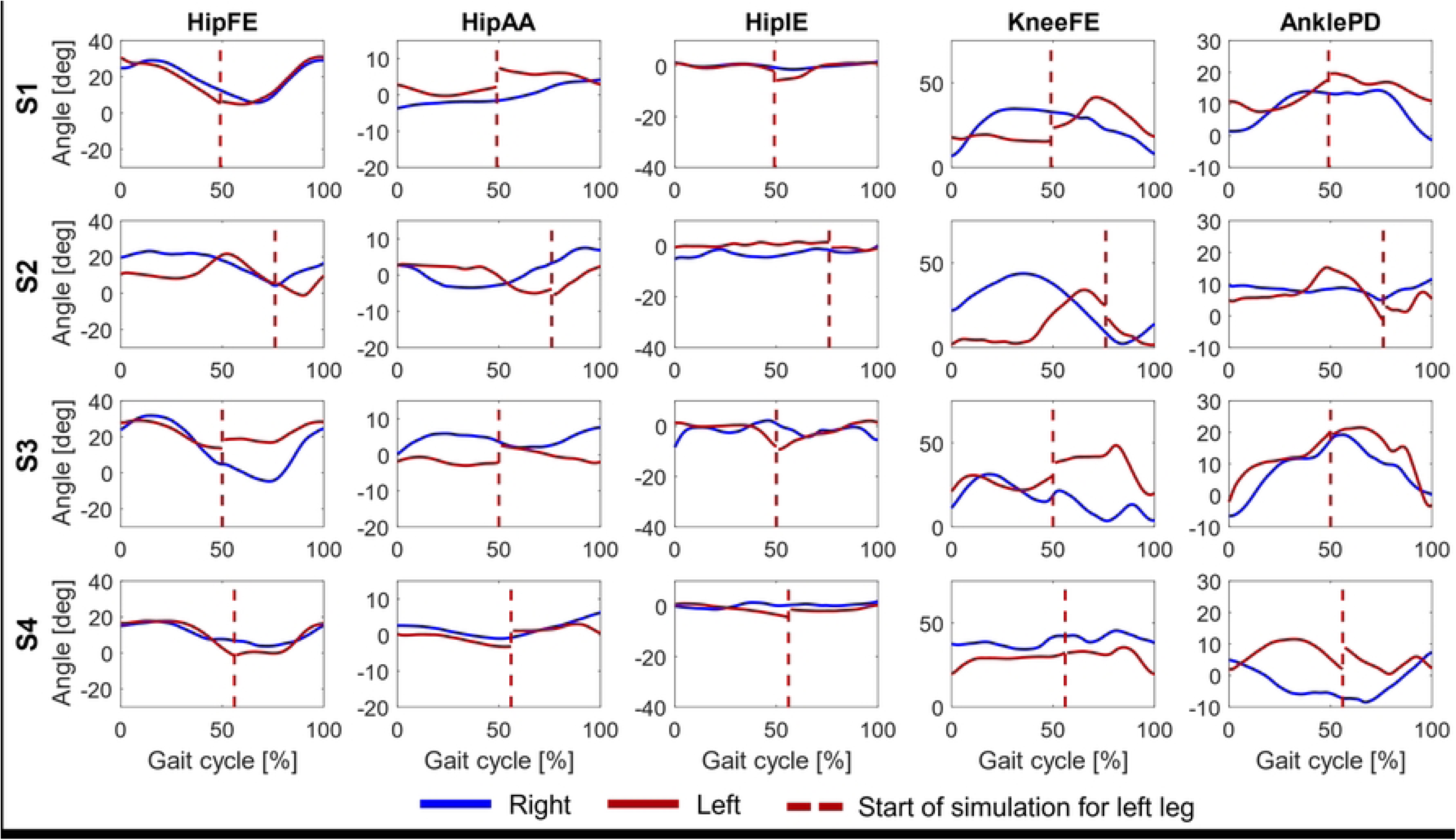
Predicted joint angles profiles. The predicted angle profiles of hipFE, hipAA, hipIE, kneeFE and anklePD of the four subjects, using FCMs. The joint angles of the left leg were rearranged to show the angle profiles between the two heel strikes of the left leg.

In the lower body kinematics, the experimental intra-subject RMSE and R^2^ values were, respectively, lower and higher than the inter-subject values of the predictive simulations with the FCMs (Fig 3). This did not always occur in the upper body kinematics, where the experimental intra-subject RMSE value of S2 and the experimental intra-subject R^2^ values of S1 and S3 were not statistically different from the predicted inter-subject variability/correlation. Furthermore, in the upper body kinematics, the mean inter-subject RMSE value of the predictions with the FCMs was similar to the experimental inter-subject value. However, there was a statistically significant difference between the predicted and experimental inter-subject RMSE/R^2^ values in the lower body kinematics. Moreover, the correlation between the experimental upper body kinematics and the corresponding predictive simulation results was statistically higher than the experimental inter-subject correlation. In contrast, the opposite occurred in the lower body, where the experimental inter-subject correlation was significantly higher. In both the lower and upper body, the inter-subject variability in the predictive simulations was significantly lower than the RMSEs between the experimental data and their corresponding predictive simulation results.

Regarding the gait metrics, there was no statistically significant difference between the experimental and predicted inter-subject variability in the circumduction and circuity index (Fig 4). However, there were statistically significant differences for the other metrics. More specifically, in the muscle co-contractions and the ROMs of the joint angles, the inter-subject variability in the predictive simulations was significantly lower than the experimental one. Additionally, except for mean trunk inclination, the differences between predicted and experimental gait metrics were significantly smaller than the experimental inter-subject variability.

According to the RMSE and R^2^ values in Table 3, the predictive simulations most accurately reproduced the upper body kinematics of its corresponding subject. In fact, when each simulation was compared against the experimental data of all subjects, none predicted the upper body kinematics of a different subject better than those of its own. Based on the RMSE values between predicted and experimental lower body kinematics, the simulation of S2 best captured the experimental data of that subject. The prediction of S4 reproduced most accurately the experimental data of S2, but also performed well for S4. In contrast, the simulation of S3 predicted more accurately the experimental lower body kinematics of the other subjects. Based on the R^2^ values, the predicted lower body kinematics of S4 correlated well with their respective experimental data, compared to the other subjects.

**Table 3.**
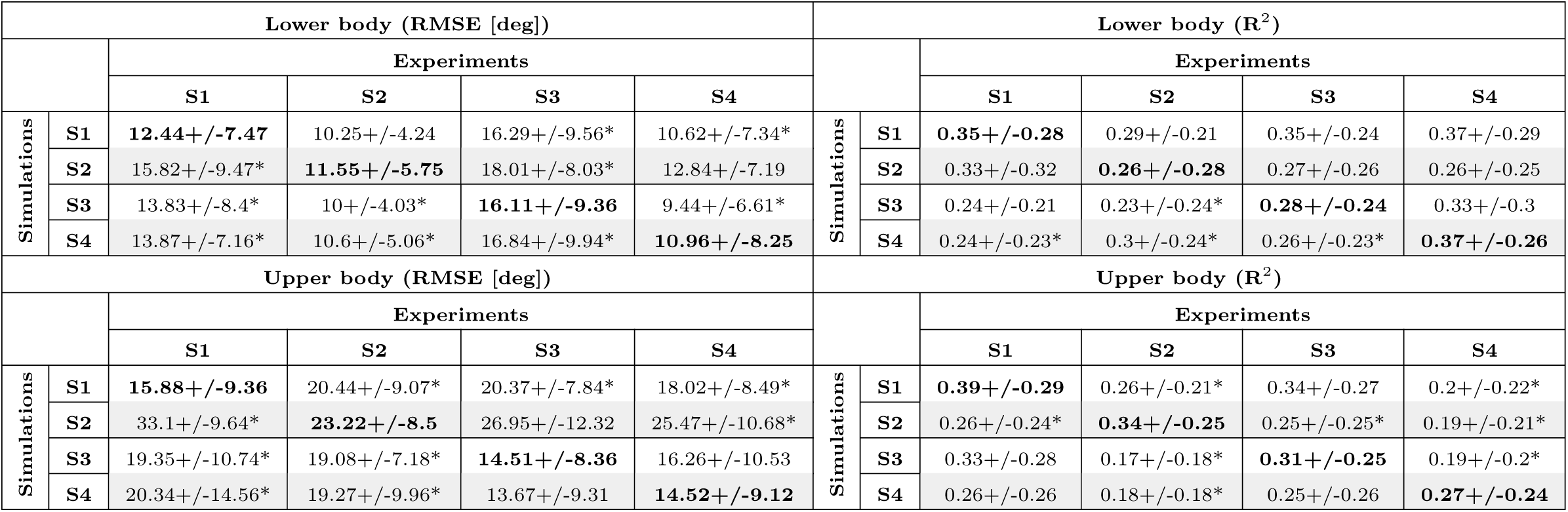
Comparison between the experimental and predicted kinematics (with FCMs). Each cell contains the mean RMSE/R^2^ value +/- its standard deviation. The bold values indicate the comparisons between the predictive simulations and the experimental measurements of the corresponding subject. In each row, the asterisks indicate a statistically significant difference with the bold value in the same row.

The simulation of S1 accurately predicted the circuity index, mean trunk inclination and upper body ROMs of that particular subject, compared to the other individuals (Table 4). In addition, the prediction of S4 reproduced more accurately the lower body ROMs of S4, compared to the other subjects. The simulation of S4 also reproduced the circuity index, mean trunk inclination and the muscles co-contractions of S4 significantly better than other two subjects. However, in some cases, the predicted metrics of a subject were statistically more similar to the experimental metrics of another subject. For example, the simulation of S2 reproduced more accurately the experimental mean trunk inclinations of S3 and S4 compared to the one of S2.

**Table 4.**
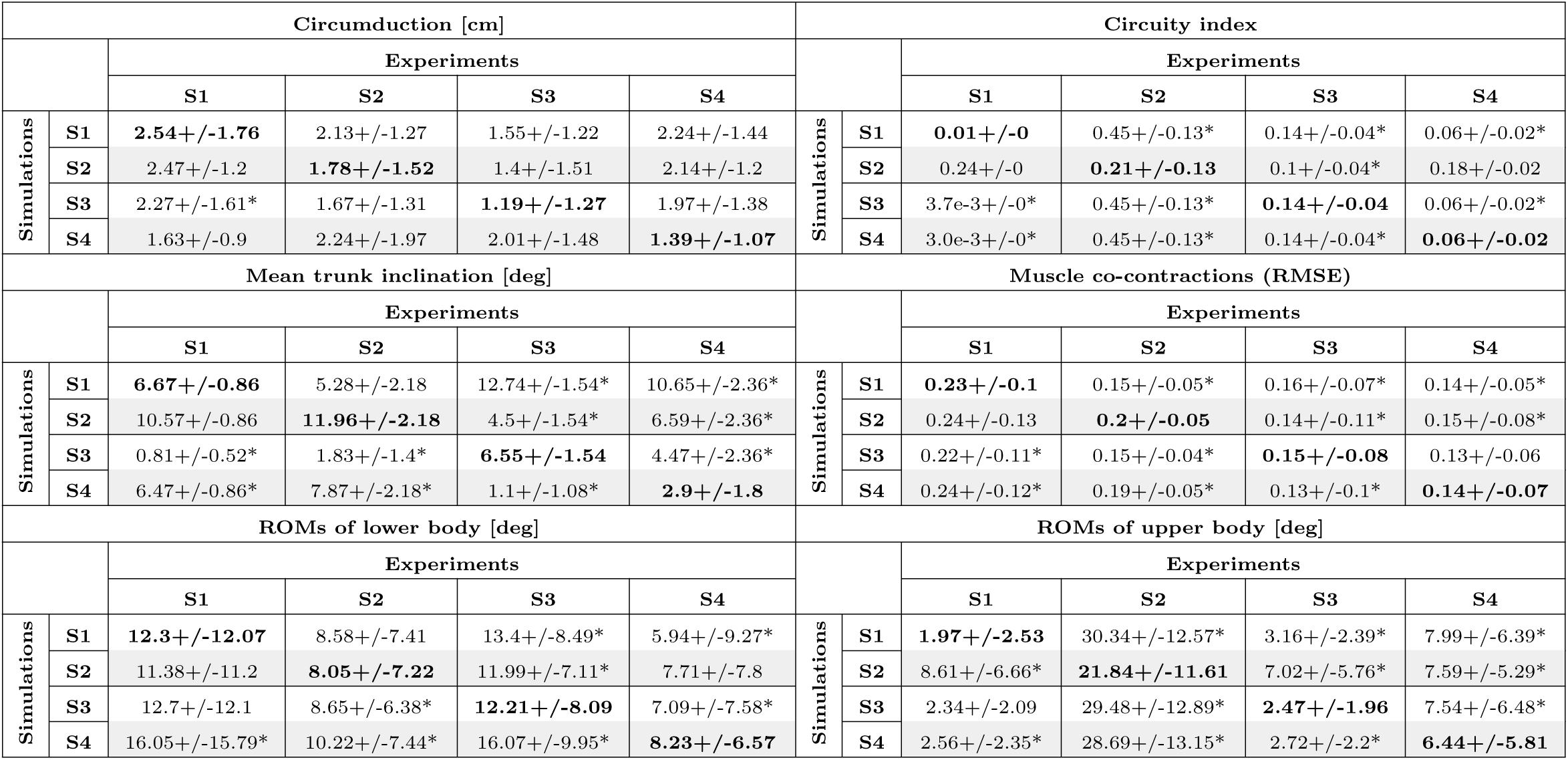
Comparison between the experimental and predicted gait metrics (with FCMs). Each cell contains the mean difference/RMSE value +/- its standard deviation. The bold values indicate the comparisons between the predictive simulations and the experimental measurements of the corresponding subject. In each row, the asterisks indicate a statistically significant difference with the bold value in the same row.

### Comparison between models

As shown in Fig 6a, the right hipFE, kneeFE and anklePD angles predicted with FCMs and FCMs-*τ_a_* were similar to each other, compared to the predictions obtained with NSMs. However, in S4, the joint angles predictions of all three models showed strong similarities. In S2, the predicted joint angles of the left leg changed significantly between simulations with FCMs and FCMs-*τ_a_* (see S3 Fig in S1 File). In addition, compared to NSMs, both FCMs and FCMs-*τ_a_* yielded statistically lower RMSE values between the predicted and experimental lower body kinematics (Fig 6b). However, based on the R^2^ values, there was no statistical difference between NSMs and the other models. In contrast, the lower body kinematics predicted using the FCMs correlated more accurately to the experimental values compared to the FCMs-*τ_a_*. In the prediction of the upper body kinematics, there was no statistically significant differences between any of the models. Moreover, the difference between the predicted and experimental circumduction values was significantly lower when using FCMs compared to NSMs (see S4 Fig in S1 File). However, for the other gait metrics, there was no statistically significant difference between the models in their prediction accuracy.

**Fig 6.**
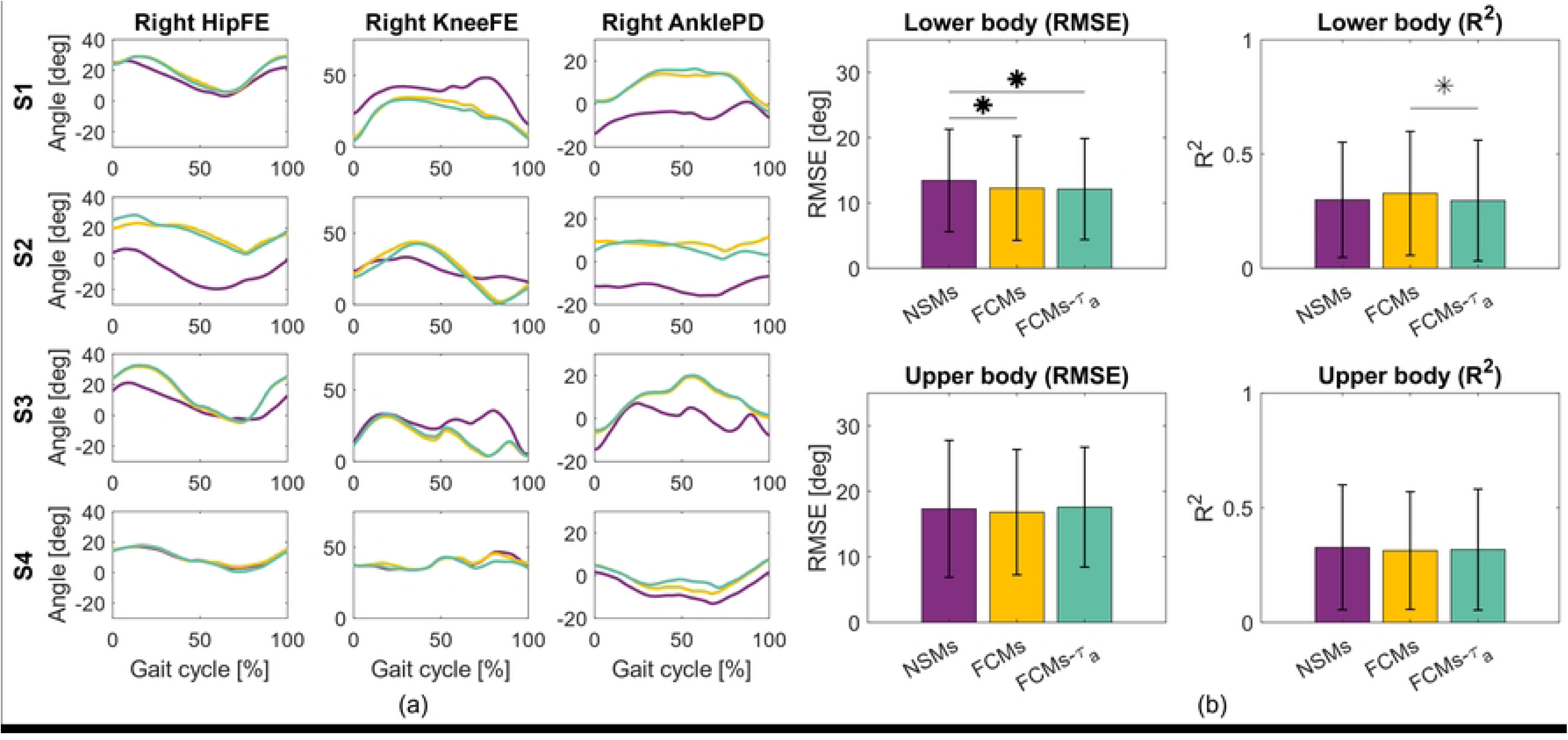
Comparison between the different types of models. (a) The predicted angle profiles of the right hipFE, kneeFE and anklePD of the four subjects, using NSMs, FCMs and FCMs-*τ_a_*. (b) The comparison between the experimental and predicted kinematics using NSMs, FCMs and FCMs-*τ_a_*. The bar plots indicate the mean difference/RMSE value, whereas the error bars indicate the respective standard deviations. Asterisks indicate statistically significant difference between two samples. If the asterisk is bold, the statistically significant difference represents an expected outcome (as specified in Table 2).

## Discussion

In this study, we evaluated the ability of our predictive simulation framework with different calibrated musculoskeletal models to reproduce and distinguish the gait characteristics of four subjects with SCI. To do so, we first analyzed the experimental data to identify the subject-specific features of the participants’ walking patterns. Then, we generated predictive simulations of walker-assisted gait with FCMs to assess if they accurately captured the distinct gaits of the four subjects. Lastly, we assessed which level of model personalization is necessary to predict subject-specific walking motions in individuals with SCI. Overall, the results suggest that our predictive simulation framework is able to reproduce distinct gait metrics of specific subjects with SCI. The predicted inter-subject variability of both the kinematics and the gait metrics is, however, generally lower than that measured experimentally. The simulations with FCMs and FCMs-*τ_a_* produced lower RMSE values between the predicted and experimental lower body kinematics compared to NSMs.

The stiff knee angle profiles predicted for S1, S3, and S4 using FCMs may be explained by several factors. The passive muscle forces of certain knee flexors (e.g., the biceps femoris long head and short head for S1) were high during stance. Since the cost function of the OCP minimized passive joint torques, further extension of the knee was discouraged from the optimization to avoid an excessive increase in passive knee torques. Inaccurate estimations of kneeFE angles caused by high passive forces were also reported in our previous study [47] and in another work [22]. In addition, since we scaled the 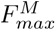 parameters of the muscles based on the LEMS results of the patients, some knee extensor muscles might have resulted too weak to counter-act the high knee flexors passive forces during stance. In fact, the knee extensors of S1’s left leg and S3’s right leg were assigned LEMS scores of 2/5 and 3/5, respectively. Moreover, as described in another predictive simulation study [30], a crouch gait with stiff knees could result as more optimal compared to an upright walk in OCPs. In fact, in that study, the crouch gait produces lower muscle activations and passive torques, which lead to a lower cost function value. In walker-assisted gait, it is even more likely for crouch gait to result as more optimal than upright walking, because the support provided by the walker reduces the ground reaction forces, thereby lowering the required joint torques and muscle activations in the lower limbs. This reasoning might also explain why the hipIE and hipAA angles remain relatively constant throughout the gait cycle for all subjects. Movements outside the sagittal plane that do not contribute to the forward progression of the model might cause higher activations, passive torques and metabolic energy expenditure, thereby increasing the cost function value.

The inter-subject variability (i.e., RMSE) of the lower body kinematics predicted using FCMs was statistically lower than in the experiments (Fig 3). This occurred mainly because, in the simulations, S1, S3 and S4 exhibited similarly stiff knees throughout the gait cycle, and the joint angles outside the sagittal plane remained nearly constant across all subjects. Moreover, the inter-subject correlation (i.e., R^2^) of the lower body kinematics predicted using FCMs was also statistically lower than in the experiments. Therefore, even though the predicted joint angles were similar in magnitude across subjects, their temporal patterns differed, resulting in lower correlations. Additionally, the predicted inter-subject correlation in the upper body was statistically different from that measured experimentally and was comparable to the experimental intra-subject correlations for S1 and S3. This likely occurred due to the lack of muscle actuation and personalization of the upper body. When predicting walker-assisted gait, the upper extremities play a more active role in ambulation. Therefore, it is important to realistically model the muscle contribution in the upper body motion, a feature that is often missing in existing musculoskeletal models for gait prediction. This need becomes even more evident when considering the predicted inter-subject variability of upper body ROMs, which are not statistically different from the experimental intra-subject variability values (Fig 4). Furthermore, all gait metrics predicted with FCMs had a comparable or statistically lower inter-subject variability than that measured experimentally, with the exception of the mean trunk inclination. In fact, in the experiments, the subjects walked with a similar trunk inclination, whereas in the simulations, S2 and S4 relied much more heavily on the walker. As a result of the high HRFs, the models of S2 and S4 leaned their entire body onto the walker. This could have not occurred in reality as the walker would move forward and the subjects could lose balance. Implementing a contact model between the ground and the walker could therefore improve the realism of the simulated assisted gait.

Even if the overall RMSE values between the experimental gaits and their corresponding predictive simulations with FCMs are not significantly lower than the overall experimental and predicted inter-subject variability (Fig 3), specific subject-to-subject comparisons are, in some cases, well reproduced in the FCMs predictions. For instance, based on the RMSE values in Table 3, the simulation of S4 most accurately captures the experimental data of S2 and S4, reflecting the experimental similarities between these two subjects (S1 Table in S1 File). Moreover, the FCMs simulation of S2 predicts the experimental lower body kinematics of S3 the least accurately, a distinction that is consistent with the experimental data. This is considered a promising result because, according to the clinical scales reported in Table 1, S3 walked considerably better than S2, and our predictive simulation framework successfully captured this inter-subject difference. Similar observations can be made when considering the R^2^ value in Table 3 and S1 Table (in S1 File). For example, the predicted upper body kinematics of S1 and S3 show significantly stronger correlations with the experimental data of both S1 and S3 than with the data of the other two subjects, mirroring the correlations observed in the experimental measurements. Since S1 and S3 have the highest WISCI-II scores among all subjects, their upper body kinematics are expected to be highly correlated, as they were able to walk with good stability and minimal movement of their arms and trunk. In fact, similar conclusions can be drawn from the ROM values in the upper bodies of S1 and S3 (Table 4 and S2 Table in S1 File). Moreover, the simulations of S1 and S4 predict most accurately the circuity indices of these two subjects, thereby reflecting their similarity in the experimental data.

In our previous work [47], we did not find a clear difference between FCMs and NSMs in predictive simulations with healthy subjects. However, in this study, both FCMs and FCMs-*τ_a_* yielded lower RMSE values between the predicted and measured lower body kinematics compared to NSMs. This observation may be the result of two main contributing factors. First, compared to our previous study, we applied tighter bounds for the 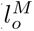 and 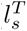 parameters in the muscle calibration OCP (i.e., +/- 10% of their baseline non-linearly scaled value). This avoided as much as possible high passive muscle forces that would prevent the model from reaching certain joint angles. Second, calibrating the musculotendon parameters through functional approaches may be critical for patients with SCI, whereas anthropometric methods could be sufficient to predict the gait for healthy individuals. In fact, the muscle properties in impaired subjects may vary more than in healthy people, and therefore, their models’ parameters cannot be simply scaled based on skeletal dimensions. The results from this work align with the findings in other predictive simulation studies involving children with cerebral palsy [30, 31]. In [30], the RMSE between the predicted and experimental kneeFE angles of a child were lower when using a FCM. When using a generic model, the simulations predicted an upright gait pattern, thus failing to reproduce the characteristic crouch gait of the child. Van Den Bosch *et al.* [31] modeled muscle contractures by reducing the 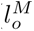 values of the Hill-type units. To do so, they placed the musculoskeletal model in the same positions of the children during passive ROM assessments and determined the 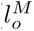 value to match a specific net joint torque value. The models with reduced 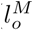 parameters achieved lower RMSE values between the predicted and measured hipFE and kneeFE angles compared to generic models. Furthermore, as shown in Fig 6a, the right joint angles predicted with FCMs and FCMs-*τ_a_* are very similar. However, the R^2^ values between the experimental and predicted lower body kinematics when using FCMs and FCMs-*τ_a_* showed a statistically significant difference (Fig 6b). This occurred because, in S2, the calibrated *τ_a_* values changed significantly compared to the baseline and, consequently, the joint angles in the left leg predicted with FCMs-*τ_a_*were substantially different than with FCMs. Therefore, our results suggest that calibrating *τ_a_* affects the predictive simulations only when the parameter varies significantly from the baseline. Based on the findings of this study, we cannot conclude whether estimating subject-specific *τ_a_* values improves gait prediction results in individuals with SCI.

This study has some limitations in relation to the modeling methods, formulation of the predictive simulations problem and number of subjects. Regarding the modeling approaches, we did not add muscle units in the upper body to avoid a significant increase in the computational time of the predictive simulations. Future studies should consider implementing muscle torque generators (MTGs) in the upper body to combine physiological realism with computational efficiency. MTGs are rotational, single muscle equivalents that contain joint angle/velocity scaling and passive elements to mimic the Hill-type muscle model behavior [67, 68]. In addition, the interaction between the hands and the walker was modeled by applying three hand reaction forces at each hand. A more realistic hands-walker and walker-ground contact model should be implemented to improve the realism of the whole system. Regarding the formulation of the predictive simulations problem, we bounded some key parameters, such as the stride length, swing/stance phase durations, PWS and gait speed to match the experimental values (within a tolerance). Future studies should focus on performing gait predictions capable of accurately reproducing these key gait metrics without constraining them. Moreover, the weights of the cost function terms were selected through trial and error. Further work should be carried out to use multi-objective optimization to explore the entire Pareto-optimal front of the control objectives and identify their optimal weights [69]. Lastly, more subjects are required to achieve more conclusive and generalizable results. This is especially important when performing predictive simulations with subjects with neuromuscular diseases or traumatic injuries because gait patterns might vary significantly between each individual. Despite these limitations, this study provides encouraging results supporting the potential of predictive simulations for future clinical decision-making processes.

## Conclusion

In this study, we assessed if our predictive simulation framework could reproduce and distinguish the subject-specific gait patterns of four individuals with SCI. To this end, we first analyzed the experimental data to characterize the gait of each subject, then we evaluated if the predictive simulations using functionally-calibrated models generated the distinct walker-assisted gait features of the four patients, and lastly, we examined how different levels of musculoskeletal model personalization affected the predictive simulation results. Our results suggest that the predictive simulations with functionally-calibrated models were able to reproduce some subject-specific gait metrics for the four individuals with SCI. Moreover, when comparing subjects pairwise, in certain instances, the predictive simulations captured the similarities or discrepancies in kinematics and gait metrics between two individuals. However, the predicted inter-subject variability of both the kinematics and the gait metrics was generally statistically lower than the experimental one. Additionally, functionally-calibrated models produced more accurate lower body kinematics results compared to non-linearly scaled models. Future work will focus on increasing the realism of musculoskeletal models (e.g., by increasing the complexity of the hand-walker contact model), improving the formulation of the predictive simulation framework (e.g., by using multi-objective optimization to determine the optimal weights of the control objectives), and including more subjects for attaining more conclusive and generalizable results.

## Supporting information

**S1 File. Supplementary material.** A PDF file including more details regarding the predictive simulation framework, along with additional figures and tables.

**S2 File. Experimental data and simulation results.**

## Acknowledgments

We would like to thank the Biomechanics and Technical Aids Department of the Hospital Nacional de Parapléjicos (in Toledo, Spain) for recruiting the patients included in this study and for conducting the experiments.

